# Characterization of erythroferrone oligomerization and its impact on BMP antagonism

**DOI:** 10.1101/2023.09.01.555965

**Authors:** Eleanor M Mast, Edmund A E Leach, Thomas B Thompson

## Abstract

Hepcidin, a peptide hormone that negatively regulates iron metabolism, is expressed by bone morphogenetic protein (BMP) signaling. Erythroferrone (ERFE) is an extracellular protein that binds and inhibits BMP ligands, thus positively regulating iron import by indirectly suppressing hepcidin. This allows for rapid erythrocyte regeneration after blood loss. ERFE belongs to the C1Q/TNF related protein (CTRP) family and is suggested to adopt multiple oligomeric forms: a trimer, a hexamer, and a high molecular weight species. The molecular basis for how ERFE binds BMP ligands and how the different oligomeric states impact BMP inhibition are poorly understood. In this study, we demonstrated that ERFE activity is dependent on the presence of stable dimeric or trimeric ERFE, and that larger species are dispensable for BMP inhibition. Additionally, we used an **in-silico** approach to identify a helix, termed the ligand binding domain (LBD), that was predicted to bind BMPs and occlude the type I receptor pocket. We provide evidence that the LBD is crucial for activity through luciferase assays and surface plasmon resonance (SPR) analysis. Our findings provide new insight into how ERFE oligomerization impacts BMP inhibition, while identifying critical molecular features of ERFE essential for binding BMP ligands.

## Introduction

Erythroferrone (ERFE) is the primary erythroid regulator of iron metabolism.^1^ When iron overload is sensed, bone morphogenetic protein (BMP)-induced hepcidin transcription occurs in hepatocytes, which negatively regulates gut iron import.^2–4^ After a blood loss event, erythroid precursor-produced ERFE directly antagonizes BMP ligands.^1,5,6^ By turning off this negative regulatory mechanism, ERFE positively regulates iron import, allowing for rapid regeneration of erythrocytes. ERFE contributes to iron overload in a mouse model of β-thalassemia which, like human patients, experience pathogenic erythropoiesis and iron overload.^7–9^ Both liver and spleen iron overload in these mice was partially alleviated after treatment with an ERFE neutralizing antibody or genetically by crossing β-thalassemic mice with ERFE^-/-^ mice.^7,8^ Thus, a better understanding of ERFE and its mechanism of action may lead to a better comprehension of the role of ERFE in pathophysiological states of iron regulation.

ERFE is part of the 16-member C1Q/TNF related protein (CTRP) family. While ERFE is the only member known to antagonize BMP ligands, it shares the family’s characteristic form (Fig**1.A**).^10^ CTRPs are named for their C-terminal globular C1Q domain^10^; numerous crystal structures of C1Q domains depict a TNF-α-like trimer held together by noncovalent interactions.^11–13^ CTRPs also possess an N-terminal unstructured domain (USD) containing unpaired cysteines and a proline/glycine-rich collagen-like repeat (CLR) which, like collagen, is predicted to form a triple helix.^10,13–15^ Well-characterized family members are known to trimerize using their C1Q and CLR and form higher order multimers by linking trimeric units together with intermolecular disulfide bonds.^16–18^ While published data suggest ERFE may oligomerize similarly^19,20^, the link between ERFE oligomerization and activity remains unexplored.

**Figure 1:**
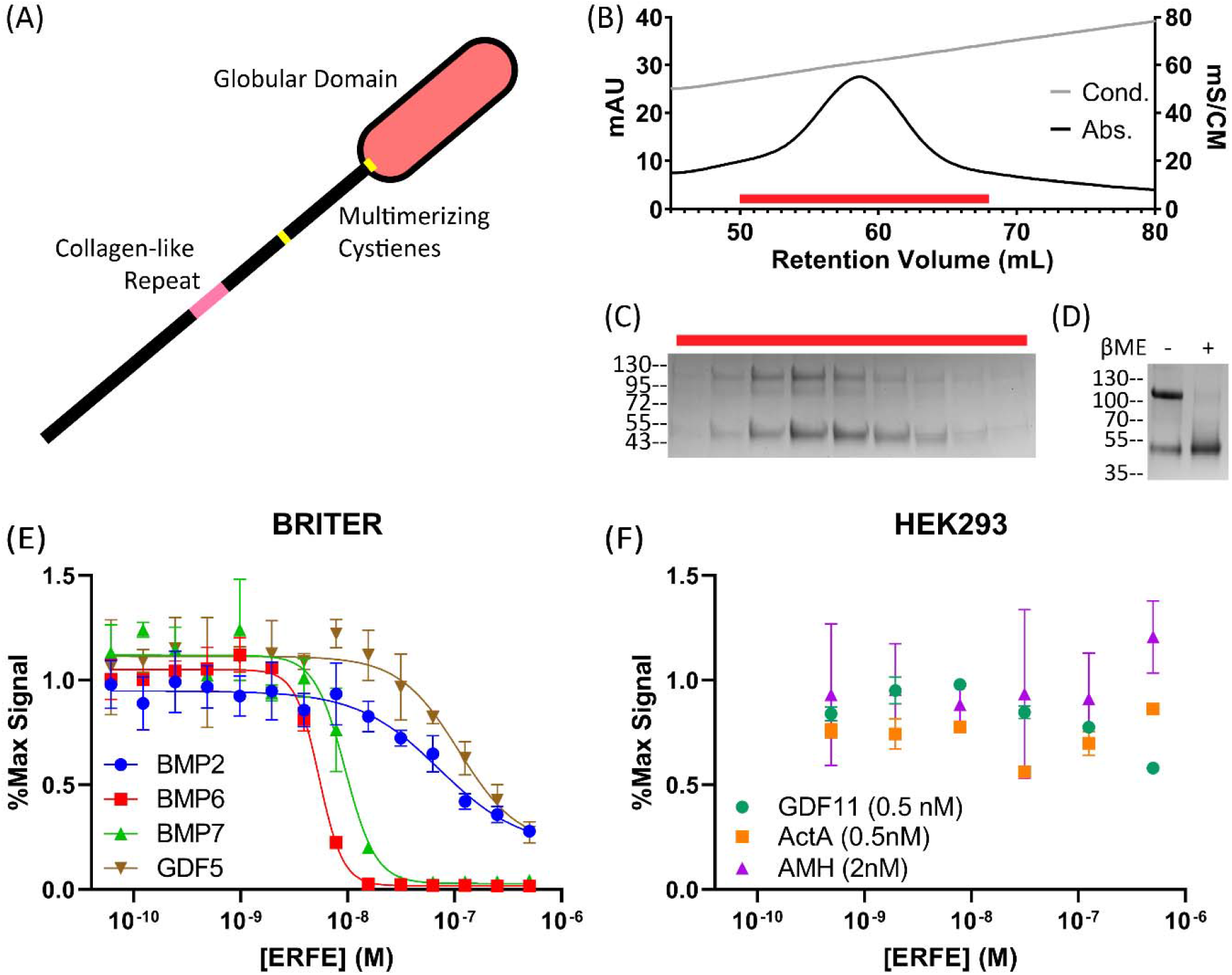
Purified ERFE is a BMP-specific antagonist. (**A**) Schematic of with the C1q domain shown as an oval. The USD contains the collagen-like repeat and cysteine predicted to aid in oligomerization. (**B**) WT ERFE purified using heparin affinity chromatography and eluted with a salt gradient. (**C**) Peak fractions corresponding to the red bar were analyzed via SDS-PAGE. (**D**) Purified ERFE analyzed under reducing and non-reducing conditions. ERFE contains a disulfide-linked dimer and monomer component. (**E**,**F**) ERFE inhibition was measured via BMP responsive luciferase assay using a BRE promoter. ERFE was titrated against 2.5nM BMPs in BRITER cells (n=3, representative curve) and different concentrations of other ligands (n=2, representative curve) in HEK293 cells, all starting at 500nM ERFE. Statistical analysis is detailed in materials and methods and results are in Table 1. SD of 3 points within a single technical replicate shown with error bars.

ERFE is known to bind and inhibit Bone Morphogenetic Proteins (BMPs), which are the largest subfamily of the transforming growth factor-β (TGF-β) family of signaling ligands.^6^ Like all TGF-β ligands, BMPs signal by binding to a pair of type I receptors and a pair of type II receptors, leading to receptor activation and SMAD 1/5/9-dependent changes in DNA transcription.^21,22^ BMPs are heavily regulated by a number of structurally diverse extracellular antagonists which tightly bind to BMPs, occluding their receptor binding sites.^21,23,24^ These antagonists bind with varying specificity across the subfamily of BMP ligands and are in large part driven by distinct binding mechanisms. These mechanisms include differences in binding stoichiometry – chordin (monomer) and noggin (dimer) both bind BMP ligands in a 1:1 ratio, whereas gremlin-2 (dimer) binds in a 2:1 ratio.^24–26^ ERFE, a trimeric inhibitor, represents a potentially unique and uncharacterized binding modality found within BMP antagonists.

In this study, we demonstrated that ERFE potently inhibited BMP subfamily members BMP6 and 7, and inhibited BMP2 and GDF5 to a lesser degree. We determined the functional inhibitory unit of ERFE is a trimer, as oligomeric states above a trimer did not differ in BMP6 inhibition and states below trimer were less efficacious. Finally, we used AlphaFold modeling and cross-species conservation analysis to identify a highly conserved helix in ERFE’s USD that was predicted to bind in the same location as type I receptors. By utilizing site-directed mutagenesis and surface plasmon resonance (SPR), we provide evidence that a conserved helix within the USD, which we termed the ligand binding domain (LBD), is critical for BMP antagonism by inhibiting BMP:type I receptor interaction.

## Results

### ERFE is a BMP-specific antagonist

To produce recombinant ERFE, we transiently expressed an N-terminally flag-tagged construct (provided by Dr. Tomas Ganz)^27^ in Expi293 cells. Conditioned media was purified via heparin affinity chromatography by elution with a NaCl gradient (Fig**1.B**). The resulting protein was >95% pure when analyzed via SDS-PAGE and all bands were confirmed to include ERFE as shown by both anti-flag and anti-ERFE western blots (Fig**1.C**, Supporting Figure **1.A-B**). The majority of ERFE appeared primarily in 2 distinct bands with molecular weight corresponding to a monomer and a disulfide-linked dimer, as was previously reported^19^ (Fig**1.D**). The inhibitory activity of purified ERFE was tested with a luciferase assay in a BMP-responsive cell line (Fig**1.E**, Table **1**).^28^ When tested against 2.5 nM BMP6, the observed ERFE IC_50_ value was 5.17 nM. This shows our purified ERFE is a highly potent inhibitor.

**Table 1:** ERFE IC_50_ values when tested against an array of TGF-β ligands (n=3).

|  | IC<sub>50</sub> (M) |
| --- | --- |
| BMP2 (2.5nM) | $1.7 \pm 0.57 \times 10^{-7}$ |
| BMP6 (2.5nM) | $5.2 \pm 0.52 \times 10^{-9}$ |
| BMP7 (2.5nM) | $8.9 \pm 0.75 \times 10^{-9}$ |
| GDF5 (2.5nM) | $1.3 \pm 0.25 \times 10^{-7}$ |
| Activin A (0.5nM) | n/a |
| GDF11 (0.5 nM) | n/a |
| AMH (2 nM) | n/a |

As previously reported, ERFE also strongly inhibits BMP7 (IC50: 8.85 nM), a member of the same BMP subclass as BMP6. ERFE less potently inhibited BMP2 and GDF5, which represent two additional BMP subclasses. Next, we examined ERFE with ligands hitherto untested for ERFE inhibition: Activin A, GDF11, and AMH. In all cases, we observed no ERFE inhibition, even at 250-fold molar excess when tested against AMH or 1000-fold molar excess for GDF11 and Activin A (Fig**1.F**). This agrees with previous reports that ERFE is a BMP-specific inhibitor and provides the additional evidence of interactions with the GDF5/6/7 subclass of BMPs.

### Collagen-like repeat- and cysteine-mediated oligomerization is dispensable for ERFE inhibition of BMP signaling

Present studies suggest that ERFE trimerizes and, through inter-trimer linkages, forms a hexamer and a high molecular weight oligomer of an unspecified size (Fig**2.A**).^19,20^ Previous studies relied on analyzing unpurified ERFE via blue native-PAGE (BN-PAGE) and detection by western blot, which yielded indistinct bands that varied between studies.^19,20^ To determine the oligomeric forms of ERFE in a more native state, we used sedimentation velocity analytical ultracentrifugation (AUC), a robust biophysical method for protein oligomeric analysis. Inconsistent results were seen when using frozen ERFE, potentially due to variations in freezing and thawing. Thus, all AUC experiments were completed using unfrozen, freshly purified protein. As the predicted frictional ratio of each c(s) peak diverged, c(s) curve fitting was performed starting at the most frequently occurring frictional ratio. Different protein populations had different frictional ratios; therefore, it was impossible to simultaneously solve for their precise molecular weights. This led to discrepancies between expected and observed weights. Consequently, approximate molecular masses were used to assign oligomeric states in a qualitative manner. Native ERFE was revealed to be primarily trimer (∼44%, ∼106 kDa) and hexamer (∼48%, ∼181 kDa) with a small amount of a high molecular weight species (∼7%, ∼319 kDa) (Fig**2.B**). No other higher-order oligomer was observed in substantial quantities.

**Figure 2:**
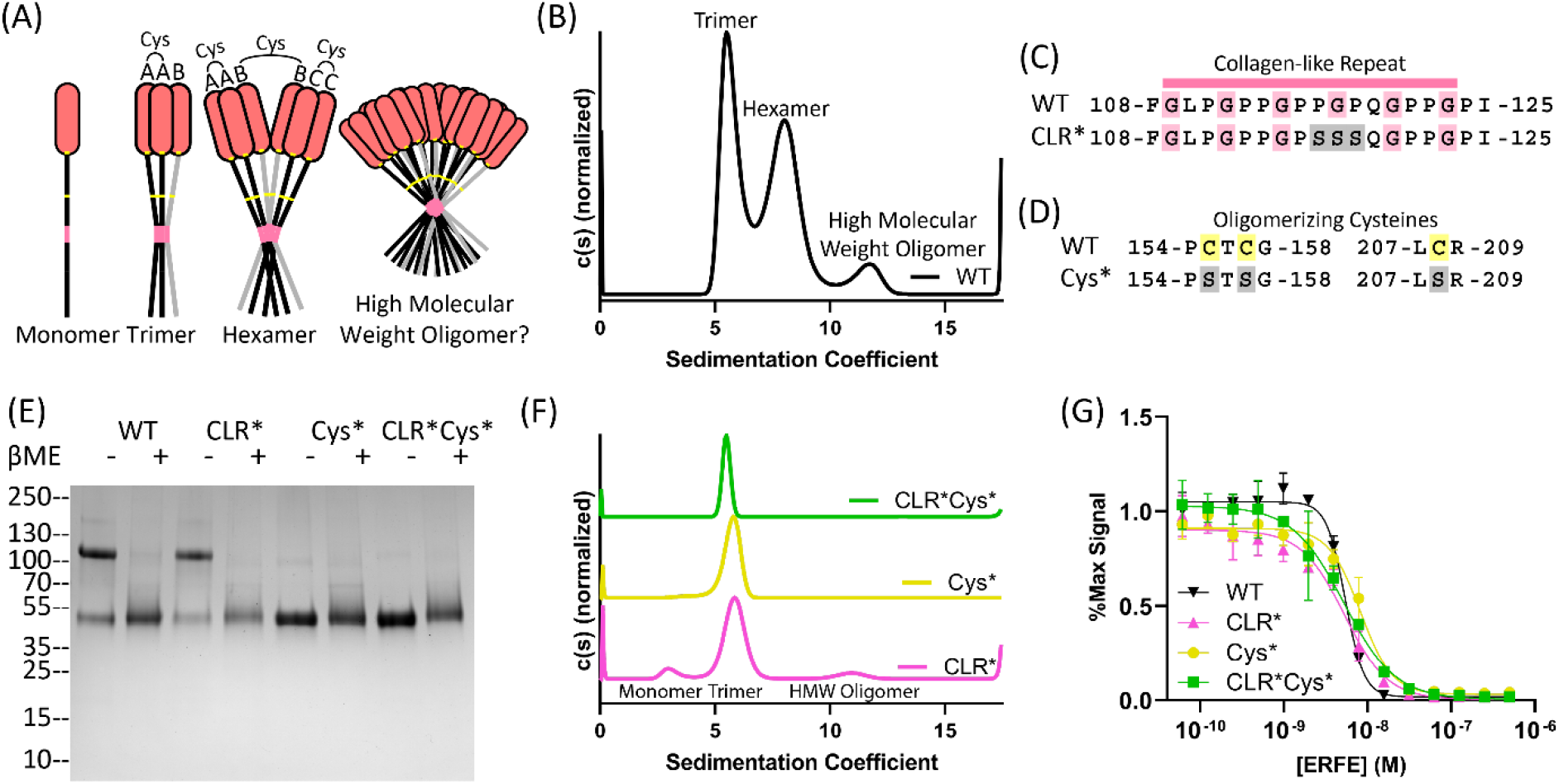
Collagen-like repeat- and cysteine-mediated oligomerization is dispensable for ERFE inhibition of BMP6. (**A**) ERFE is predicted to form higher order oligomers with a trimer as the basic unit, held together by intermolecular disulfide bonds as indicated. (**B**) Sedimentation velocity reveals that WT ERFE is predominantly trimer and hexamer, with a small fraction of high molecular weight oligomer. (**C-D**) Schematic of two mutants generated to disrupt ERFE oligomerization – CLR* disrupts the collagen-like repeat and Cys* mutates oligomerizing cysteines into serines. (**E**) 5ug Purified ERFE mutants were analyzed via SDS page under nonreducing and reducing conditions. (**F**) Sedimentation velocity of analysis of the Cys* and CLR ERFE mutants (**G**) Analysis of BMP inhibition of the mutants as compared to WT ERFE. BRITER cells were stimulated with 2.5nM BMP6 and inhibited with increasing concentrations of ERFE and mutations. Statistical analysis is detailed in materials and methods and results are in Table 2. SD of 3 points within a single technical replicate shown with error bars. (n=3, representative curve shown).

Based on the literature surrounding ERFE and other CTRPs^13,18–20^, we hypothesized that ERFE’s collagen-like repeat (CLR) stabilized trimeric ERFE and cysteines enabled inter-trimer disulfide bridges, allowing for higher order covalent oligomerization (Fig**2.A**). To interrogate the contributions of each component to oligomerization and BMP inhibition, we created two mutant forms of ERFE. First, we mutated the center PGP in the CLR to SSS (termed CLR*) (Fig**2.C**). We predicted this would disrupt formation of a collagen triple helix without causing the decreased production of ΔCLR ERFE as reported by Stewart **et al**.^**20**^ To disrupt intermolecular disulfide bonds, Cys155, 157, and 208 – which were previously reported oligomerizing cysteines^20^ – were mutated to serines (termed Cys*) (Fig**2.D**). Both variations were combined to create a 3^rd^ mutant (CLR*Cys*). After production and purification, we analyzed the ERFE mutant via SDS-PAGE. We observed that CLR* had a similar ratio of disulfide linked dimer to WT ERFE, suggesting this mutation did not affect intermolecular disulfide bonds. As expected, Cys* and CLR*Cys* appeared as a single species in nonreducing conditions (Fig**2.E**).

Intriguingly, as determined by sedimentation velocity, all ERFE mutants were deficient in oligomerization above a trimer (Fig**2.F**). However, all mutants formed trimers, suggesting the C1Q domain is the main driver in stabilizing trimeric ERFE. As expected, Cys* and CLR*Cys* do not form higher order oligomers, as they lack the requisite cysteines. Interestingly, analysis of CLR* ERFE by SDS-PAGE reveals a mixture of disulfide-linked dimer and monomer, yet sedimentation velocity analysis reveals minimal oligomerization above a trimer. This suggests that both the CLR and intermolecular disulfide bridges play a role in coordinating ERFE oligomerization.

Previous studies have shown that adiponectin, a well-studied CTRP family member, exhibits different activity and functions based on differences in its oligomeric states.^13,18^ Thus, we wanted to determine if reducing the higher-order oligomers of ERFE had an impact on BMP inhibition. As such, we tested the activity of CLR*, Cys*, and CLR*Cys* ERFE using a BMP inhibition luciferase assay (Fig**2.G**, Table **2**). The results show that all three mutants had similar IC_50_ values towards BMP6 inhibition when compared to WT ERFE. This suggests that ERFE oligomerization above a trimer is dispensable for BMP inhibition.

**Table 2:** ERFE oligomer-deficient mutant IC_50_ values for BMP6 in BRITER Cells.

|  | IC<sub>50</sub> (M) | Fold Decrease from WT |
| --- | --- | --- |
| CLR* | $6.1 \pm 0.12 \times 10^{-9}$ | 1.2 |
| Cys* | $8.7 \pm 0.85 \times 10^{-9}$ | 1.7 |
| CLR*Cys* | $4.4 \pm 0.75 \times 10^{-9}$ | 0.9 |
| ΔC1Q | $7.8 \pm 0.53 \times 10^{-9}$ | 1.5 |
| CLR*ΔC1Q | $2.7 \pm 0.17 \times 10^{-8}$ | 5.2 |
| Cys*ΔC1Q | $9.7 \pm 0.67 \times 10^{-8}$ | 19 |
| CLR*Cys*ΔC1Q | $2.9 \pm 0.11 \times 10^{-7}$ | 56 |

### Ablation of ERFE trimerization decreases inhibition of BMP6

Given its similarity to other CTRPs, ERFE trimerization has been proposed to be driven in part by its C1Q domain. Thus, in order to test if ERFE trimerization is important for BMP inhibition, we hypothesized that truncating the C1Q domain of ERFE would allow for the isolation and characterization of a monomeric form of ERFE. Here, we introduced a stop codon at C208, removing the entire C1Q domain (ΔC1Q) (Fig**3.A**). The ΔC1Q mutant was combined with the previous mutant schemata (CLR* and Cys*) to generate CLR*ΔC1Q, Cys*ΔC1Q, and CLR*Cys*ΔC1Q ERFE. Each of these 4 mutants was expressed and purified in Expi293 cells. Purity was assessed via SDS-PAGE, with bands migrating at the expected mass for all constructs (Supporting figure **2.A-D**). Both ΔC1Q and CLR*ΔC1Q were composed of disulfide-linked dimer and monomer, while Cys*ΔC1Q and CLR*Cys*ΔC1Q were monomeric.

**Figure 3:**
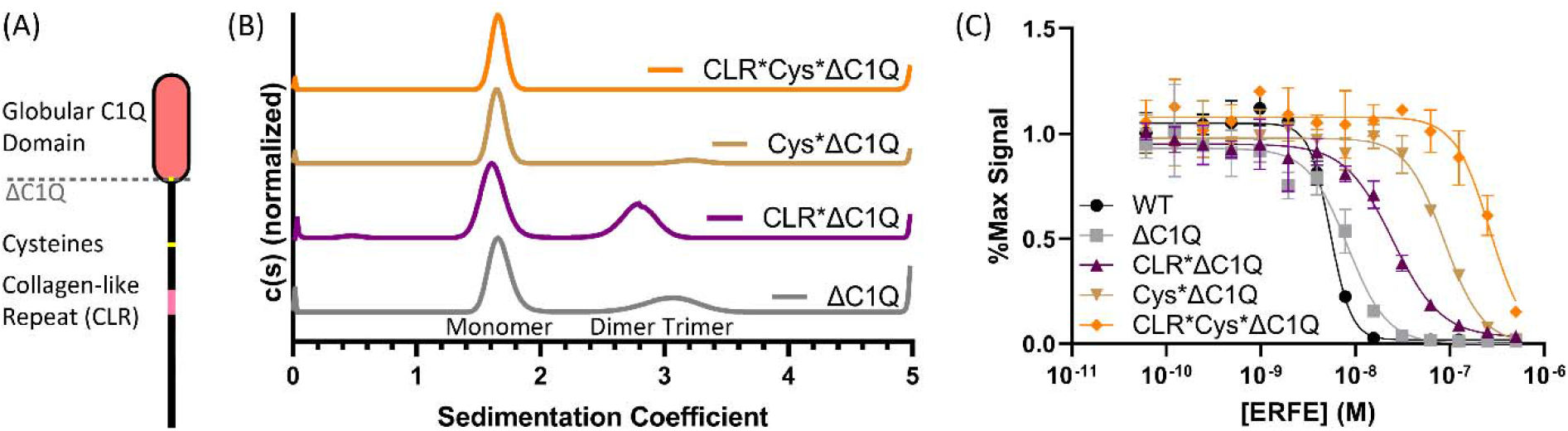
Ablation of ERFE trimerization decreases inhibition of BMP6. (**A**) Removal of ERFE’s C1Q domain to generate the ΔC1Q mutant. (**B**) Sedimentation velocity shows that ΔC1Q ERFE had reduced levels of trimer, CLR*ΔC1Q formed only monomer and dimer, dimer-deficient Cys*ΔC1Q exhibited minimal trimerization, and CLR*Cys*ΔC1Q was entirely monomeric. (**C**) BMP reporter assay to measure BMP6 (2.5nM) inhibition in BRITTER cells of the ERFE mutations. Statistical analysis is detailed in materials and methods and results are in Table 2. SD of 3 points within a single technical replicate shown with error bars. (n=3, representative curve shown).

The 4 mutants with the C1Q domain removed were analyzed via sedimentation velocity (Fig**3.B**) and molecular mass determination was performed as previously mentioned above. Interestingly, ΔC1Q did not completely disrupt trimer formation, and presented as ∼67% monomer (∼23 kDa) and ∼32% trimer (∼57 kDa). The residual trimer was removed by also disrupting the CLR, as CLR*ΔC1Q was composed of ∼61% monomer (∼19.2 kDa) and ∼38% dimer (∼43.3 kDa) with no trimer present. Surprisingly, Cys*ΔC1Q was predominantly monomer (∼20.7 kDa) with an exceedingly low amount of trimer (<8%, ∼61.6 kDa). No dimer was present, due to the lack of intermolecular disulfide bonds. Finally, CLR*Cys*ΔC1Q was entirely monomeric. These data suggest that the C1Q, CLR, and cysteines all contribute to stability of the trimer form of ERFE.

Next, we wanted to examine whether complete ablation of noncovalently associated trimer (CLR*ΔC1Q), disulfide linked dimer (Cys*ΔC1Q), or reduction to monomer (CLR*Cys*ΔC1Q) would change ERFE inhibition of BMP signaling activity (Fig**3.C**, Table **2**). Intriguingly, despite a reduction in the proportion of trimer, ΔC1Q had similar activity to WT ERFE. Ablation of trimeric ERFE, leaving monomer and dimer alone (CLR*ΔC1Q), decreased activity by 5-fold. Cys*ΔC1Q, which was predominantly monomer with a small quantity of trimer, was 10-fold less active than WT. Entirely monomeric CLR*Cys*ΔC1Q was the least active and had >50-fold less activity than WT ERFE. Taken together, these data suggest ERFE inhibition is driven by stable trimeric or dimeric ERFE.

### Identification and Mutagenesis of a Ligand Binding Domain in ERFE

Removal of the C1Q domain had little impact on BMP inhibition, supporting previously published data implicating ERFE’s unstructured domain (USD) as the main inhibitory component.^8,29^ In order to gain insight into potential binding mechanisms that enable the USD to bind and inhibit BMP ligands, we used a combination of evolutionary analysis coupled with molecular modeling. First, we used ConSurf to identify which segments in ERFE’s USD are evolutionarily conserved.^30,31^ We identified 2 conserved helices (H1 and H2) on either side of the CLR (Fig**4.A**); the sequence for the more highly conserved N-terminal helix (H1) and less conserved C-terminal helix, (H2) are shown (Fig**4.B**). Next, we used AlphaFold Multimer^32,33^ to model ERFE’s USD with a BMP6 dimer. Strikingly, in all 5 relaxed models generated, H1 was consistently placed in the BMP6 type I receptor site, while H2 was never consistently placed (Supporting figure **3.A**).

**Figure 4:**
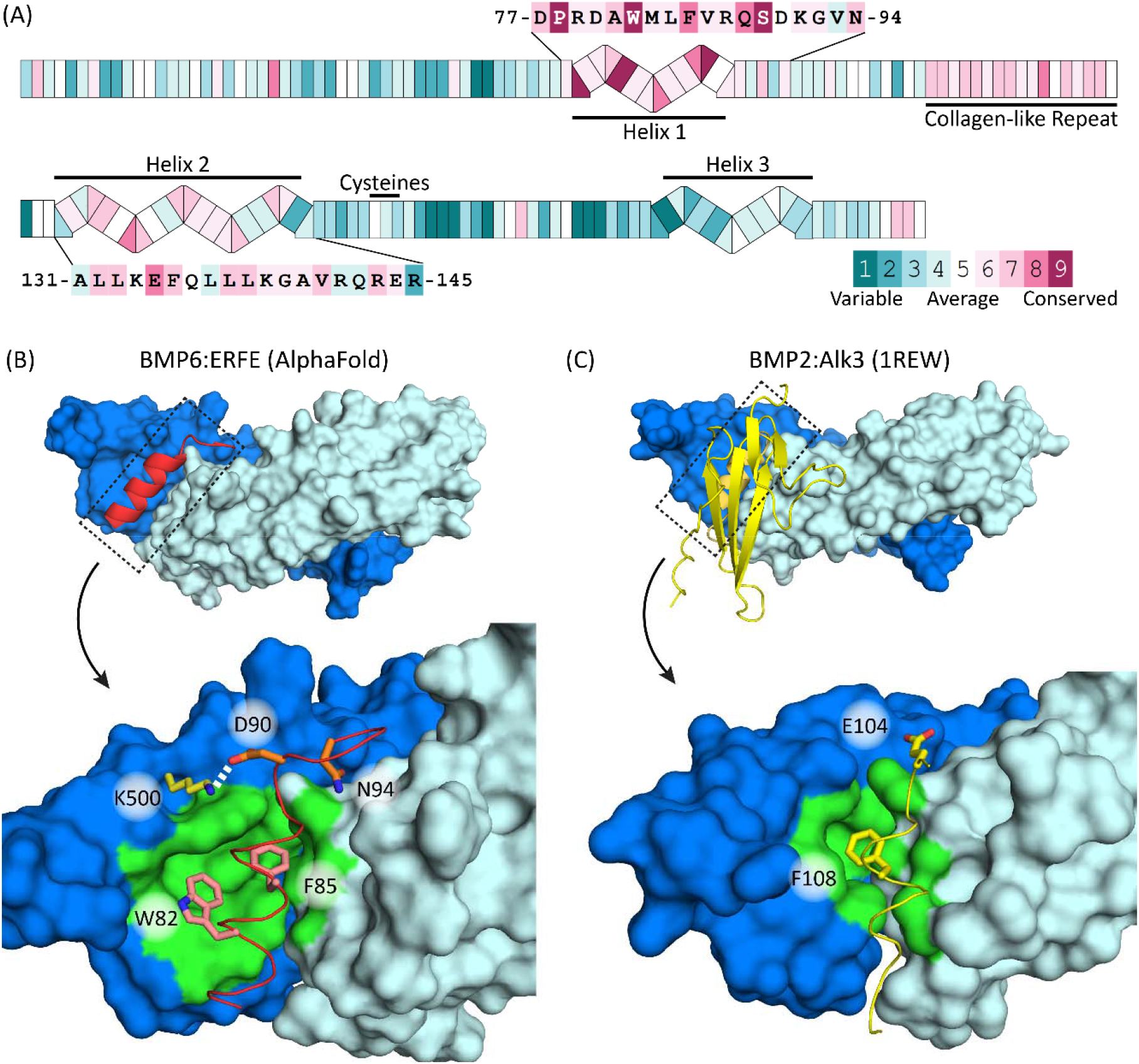
Cross-species conservation and AlphaFold molecular modeling predicted a highly conserved helix interacts with BMP ligands. (**A**) ConSurf analysis of ERFE USD. Two conserved helices (H1, H2) are labeled and sequences shown. Secondary structure predictions were derived from the publicly available AlphaFold structure database.^54^ (**B**) AlphaFold multimer modeling depicting only the H1 of ERFE bound to BMP6 (chain A and B are colored dark and light blue respectively), (lower) close-up view of H1 interactions with BMP6 showing W82, F85, D90 and N94 of ERFE interacting with the type I pocket of BMP6. K500 of BMP6 chain A forms a hydrogen bond with D90 of ERFE. (**D**) type I receptor:BMP interactions. For both C and D: bulky, hydrophobic residues are placed into the hydrophobic pocket to form a”knob-in-hole” motif and residues within 5Å of W82/F85 (ERFE) and F108 (Alk3) are colored green.

Upon closer examination, the H1:BMP6 interface buries approximately 750 Å^2^ on each component, highlighted by two conserved hydrophobic residues, W82 and F85. The predicted H1 interactions with BMP6 resembled that of other ligand binding partners (such as receptors, co-receptors, and antagonists), which increased our confidence in the model. For example, the hydrophobic interactions mirror that of type I receptor binding to BMP ligands (BMP2:ALK3 complex, PDB:1REW) (Fig**4.C-D**).^34^ Here, both utilize the common”knob-in-hole” binding motif where a bulky hydrophobic residue on either the H1 (F85) or type I receptors is inserted into a hydrophobic pocket on the ligand surface.^22,35^ Intriguingly, AlphaFold also predicted ERFE W82 interacts with two conserved tryptophans located in the concave curvature of the ligand, along with other hydrophobic residues in the ligand pocket. This interaction extends the hydrophobic interface beyond what is seen in receptor knob-in-hole interactions, broadening it towards the tip of the ligand and burying a total of 270 Å^2^ for just W82 and F85. In addition to the central hydrophobic interactions, ERFE was predicted to form ancillary interactions, such as a salt bridge from D90 of ERFE to K500 located in finger 4 of BMP6 (Fig**4.C**). This interaction resembles the capping interaction seen in other BMP:Noggin interactions.^26^ Further, N94 of ERFE is predicted to interact with the main chain of the prehelix loop of BMP6 – a ligand feature important for type I receptor specificity (Fig**4.C-D**).^34^ Thus, the modeling study is in agreement with previously characterized binding interactions and supports that ERFE may directly occlude the BMP type I receptor site.^6^

Based on these results, we designed mutants to test these predicted interactions. We first removed both the H1 and H2 helices. Removal of H1, the helix predicted to interact with BMPs, ablated ERFE inhibition of BMP signaling, while removal of H2 did not impact ERFE potency (Fig**5.A**, **Table 3**). Next, we wanted to determine if single point mutations in H1 could impact BMP inhibition. We selected 4 of the residues predicted to have significant contact with BMP6 in our model and mutated them to alanine – W82A, F85A, D90A, and N94A. Again, recombinant proteins were expressed in Expi293 cells and purified to homogeneity (Supporting figure **2.E-G**). Strikingly, almost all BMP inhibitory activity was removed by mutations W82A and F85A (Fig**5.B**, **Table 3**). D90A and N94A also decreased BMP inhibition, but to a much lesser degree: roughly a 2.5- and 4-fold loss of activity, respectively. Taken together, our data supports the BMP6:ERFE binding interaction predicted by AlphaFold. As H1, the predicted binding component of ERFE’s USD, was critical for activity, we termed it the ligand binding domain (LBD).

**Figure 5:**
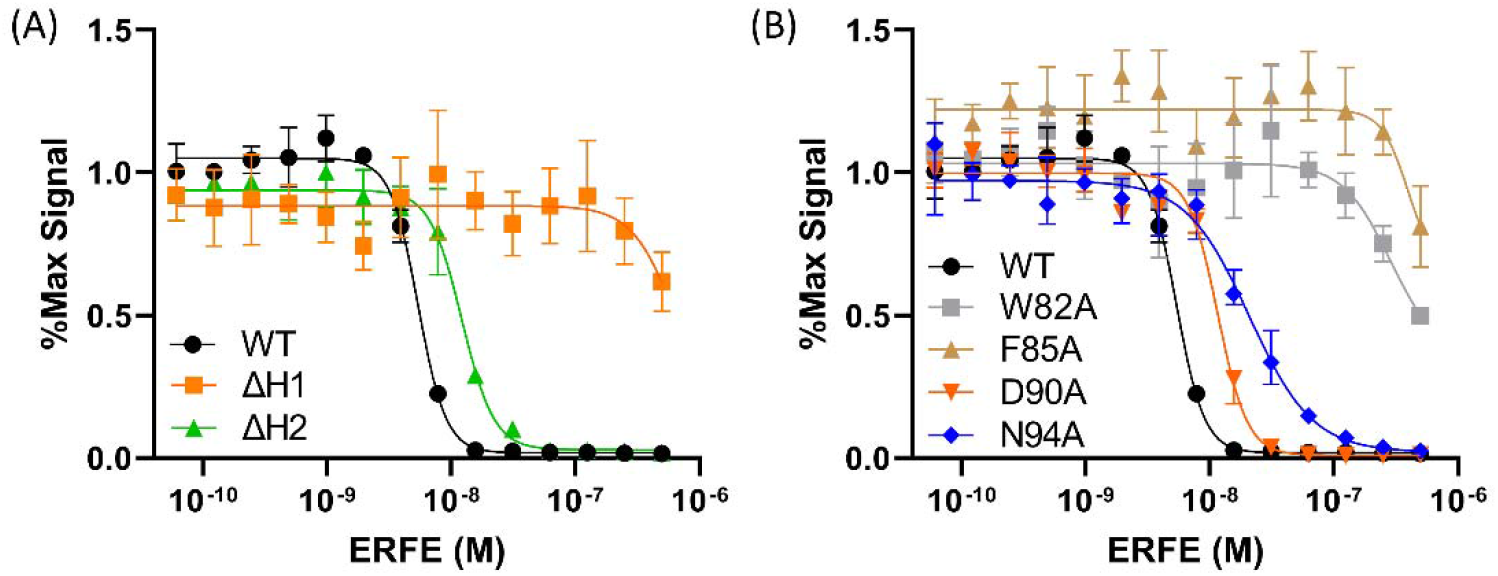
Mutation of predicted functional ERFE residues in H1 decreased inhibition of BMP6. (**A**) and (**B**) Luciferase reporter assay using BMP6 in BRITER cells. **(A**) ERFE with either H1 or H2 removed was titrated against BMP6. (**B**) Single point mutations of H1 were titrated against BMP6. Statistical analysis is detailed in materials and methods and results are in Table 3. SD of 3 points within a single technical replicate shown with error bars. (n=3, representative curve shown).

**Table 3:** ERFE Δhelix and LBD point mutant IC50 values for the inhibition BMP6 in BRITER Cells.

|  | IC<sub>50</sub> (M) | Fold Decrease from WT |
| --- | --- | --- |
| ΔH1 (ΔLBD) | n/a | n/a |
| ΔH2 | $9.1 \pm 2.6 \times 10^{-9}$ | 1.8 |
| W82A | n/a | n/a |
| F85A | n/a | n/a |
| D90A | $1.3 \pm 0.15 \times 10^{-8}$ | 2.5 |
| N94A | $2.1 \pm 0.26 \times 10^{-8}$ | 4.0 |

### ΔLBD and monomeric ERFE are less potent inhibitors of ALK3:BMP6 binding

Finally, we tested the binding model proposed by AlphaFold Multimer to determine if ERFE could disrupt ALK3 binding to BMP6 via a competition experiment using surface plasmon resonance (SPR) (Fig**6.A-B**). Alk3 was chosen as its affinity for BMP6 is highest among type I receptors, enabling greater sensitivity and lower ligand usage.^36^ In this experiment, a bivalent Alk3-Fc chimera was captured onto a protein A chip. Coupling density was kept low to avoid non-specific binding interactions and to reduce the bivalent BMP ligand interacting with multiple Fc molecules. To test if ERFE could block type I receptor interactions, we titrated increasing ERFE with constant BMP6. We predicted ERFE would bind to BMP6 and compete with type I receptor binding, leading to a decrease in response units (RU) (Fig**6.A-B**). BMP6 alone (red) exhibited tight binding to Alk3-Fc, consistent with Isaacs **et. al**.’s reported K_D_ value of 62.46 nM (Fig**6.C**).^37^ ERFE alone (grey) showed no non-specific binding to Alk3-FC, as expected^6^ (Fig**5.C**). Titrating WT ERFE with constant BMP6 showed a dose-depended decreased response in binding, suggesting disruption of BMP6 binding to Alk3 (Fig**6.C**). Intriguingly, in a similar experiment, ΔLBD ERFE did not have an impact on BMP binding to Alk3, suggesting this mutant is unable to occlude BMP6:Alk3 binding (Fig**6.D**). Similarly, fully monomeric CLR*Cys*ΔC1Q ERFE, which has very weak BMP6 inhibition, only slightly decreased binding, suggesting CLR*Cys*ΔC1Q only retained residual affinity towards BMP6 (Fig**6.E**). These data further support the hypothesis that the ERFE LBD binds to and occludes type I receptor binding.

**Figure 6:**
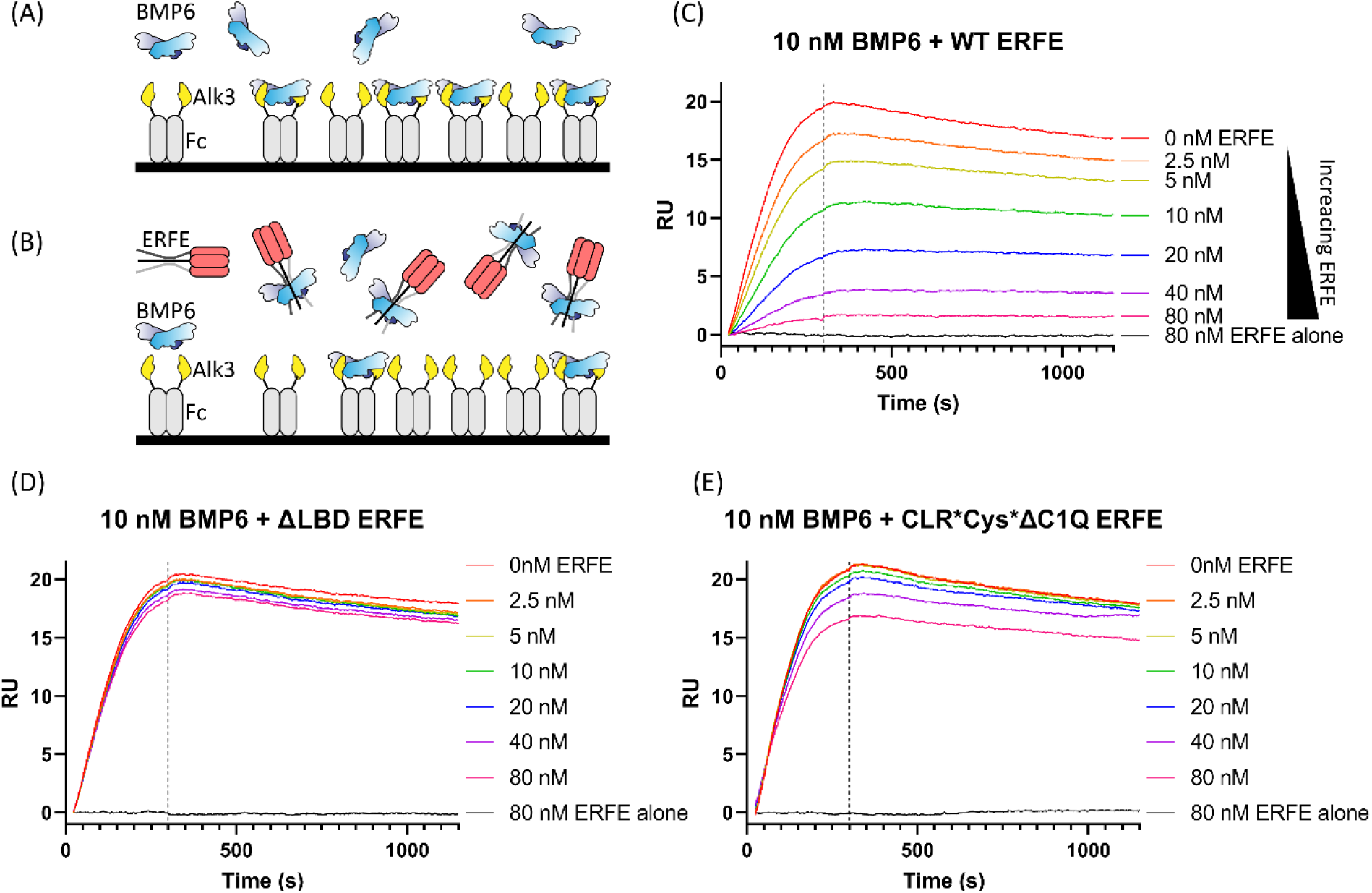
ΔLBD and monomeric ERFE are less potent inhibitors of ALK3:BMP6 binding. (**A**) SPR strategy depicting Alk3-Fc on a protein A chip capturing BMP6 ligand (**B**) By flowing over constant BMP6 and increasing the concentration of ERFE, we expected to see less BMP6:ALK3-Fc binding at higher ERFE concentrations. (**C-E**) WT ERFE or the specified mutant at the listed concentrations was precomplexed with 10nM BMP6 and flowed over coupled Alk3-Fc (n=2, representative curve shown)

## Discussion

The role of erythroferrone in iron regulation is firmly established: both its importance in rapid regeneration of erythrocytes and its contributions to iron overload. Despite this role and its potential other roles in bone growth and metabolic homeostasis^38–40^, its underlying mechanism remained understudied. Prior studies established that ERFE oligomerizes, directly binds to BMP ligands, and suggested it occluded binding to its type 1 receptors. However, neither the impact of oligomerization on activity nor details of this interaction have been examined.

To ascertain the impact of oligomerization on BMP inhibition, we first needed to better characterize WT ERFE and proceeded to define the molecular components responsible for its different states. We found that WT ERFE predominantly forms a trimer and hexamer with a small amount of high-molecular weight species via analysis with sedimentation velocity. Mutation of the cysteine residues removes the inter-molecular disulfide bonds and ablates both hexamer formation and the high MW species producing a very homogenous ERFE trimer. Disruption of the collagen-like repeat (CLR) did not alter intermolecular disulfide bond formation, but still disrupted formation of hexamer and higher-order oligomers. Thus, ERFE hexamer formation is dependent on both the cysteines and CLR.

Extending these studies to include removal of the C1Q domain helps create a better picture of ERFE oligomerization. Since all our full-length ERFE mutants were trimeric, we predicted the C1Q domain was the primary driver of ERFE trimer formation. However, ΔC1Q was only 67% monomer and the remaining 32% trimerized. The CLR stabilizes ΔC1Q, as mutation disrupted it into monomer and disulfide linked dimer. Surprisingly, mutation of the cysteines also disrupted residual trimer formation in ΔC1Q, which suggested that inter-chain disulfide bonds not only play a role in formation of higher-order oligomers, but help stabilize the CLR domain. In fact, the calculated melting temperature of ERFE’s CLR is just 9.1°C.^41^ Thus, other interactions may be required to stabilize the expected triple helix of the CLR. By extension, it is possible the lack of hexameric ERFE in full-length Cys* is due to the increased instability of the CLR, rather than the lack of intermolecular disulfide bonds. However, additional studies will be required to better characterize mechanisms of higher-order ERFE oligomerization.

Having characterized ERFE mutants that modified oligomeric states, we were better able to understand how oligomerization impacted BMP inhibition. We found that ERFE oligomerization to a hexamer and beyond was dispensable for ERFE inhibition of BMP signaling, while analysis of both WT and mutant ΔC1Q ERFE shows that dimer or trimer formation is indispensable for ERFE activity. Monomeric ERFE (CLR*Cys*ΔC1Q) was >35-fold less active than ΔC1Q. The loss in activity is likely due to a loss of avidity. Interestingly, CLR*ΔC1Q, which does not form a trimer, was only 3-fold less active than ΔC1Q. As CLR*ΔC1Q still contained disulfide linked dimer, our data suggest that, in the absence of trimer, dimeric ERFE remains active. Consistent with this, Cys*ΔC1Q, which does not form dimeric species, was just <8% trimer and was 10-fold less active than ΔC1Q. Taken together, these data support that ERFE likely functions as a trimer to inhibit BMP ligands utilizing avidity to maintain a stable inhibitory complex.

Additionally, through a combination of modeling and mutational analysis, we identified a helix in the N-terminus of the unstructured domain that binds to the type I receptor binding pocket of the ligand. As BMP ligands only exist in the body as disulfide-linked dimers, they possess two type I receptor sites.^21,22^ Trimeric ERFE would contain 3 LBDs, potentially enabling the avidity effects suggested by our ΔC1Q mutant analysis. From these data, we proposed the following model (Fig**7**). Trimeric ERFE, held together by a C1Q domain, an intra-trimer disulfide bond, and a collagen-like repeat, binds and inhibits BMP ligands. This downregulates hepcidin induction and upregulates iron import. ERFE does this by occluding BMP ligands’ type I receptor sites through direct binding with its LBDs, which forms a non-signaling ERFE:BMP complex. Furthermore, this model suggests that trimeric ERFE binds dimeric BMP ligands, potentially leaving an unpaired LBD. Two trimeric ERFE:BMP complexes with unpaired LBDs could potentially pair with a second BMP molecule to form a larger inhibitory complex (Fig**7**). This model would predict that a hexamer of ERFE would be stabilized by binding three BMP ligands. However, support for this model is lacking as attempts to characterize an ERFE:BMP complex are complicated by solubility issues.

**Figure 7:**
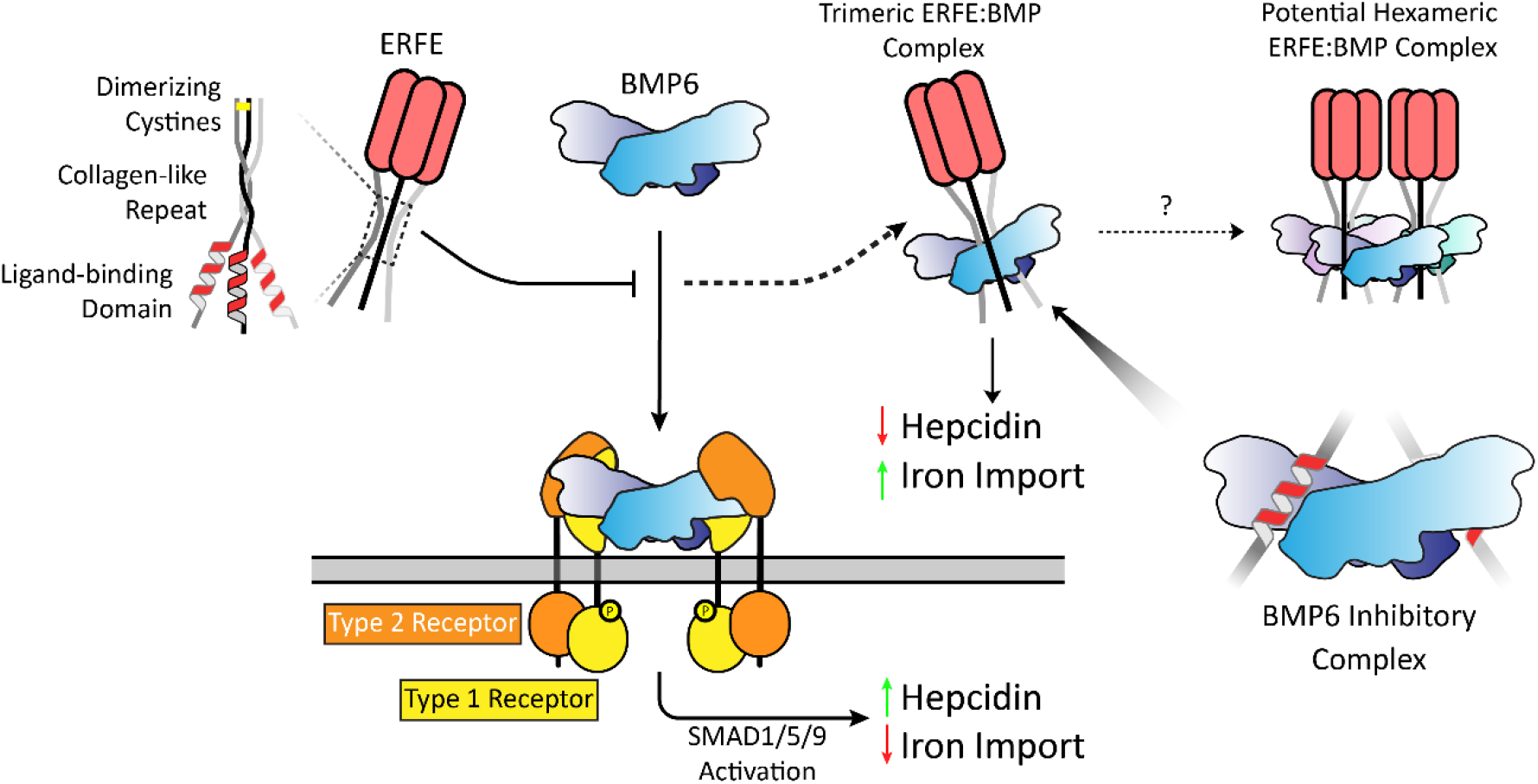
Model for iron regulation by ERFE inhibition of BMP. The active unit of ERFE is a trimer stabilized by a C1Q domain, inter-chain disulfide bonds, and CLR. When the body senses iron overload, BMP6 is produced, which induces hepcidin transcription and downregulates iron import. After blood loss, trimeric ERFE negatively regulates this pathway by directly binding and inhibiting BMP6, leading to decreased hepcidin and increased iron import. The BMP6:ERFE interaction is driven by ERFE’s conserved BMP-binding helix or ligand binding domain (LBD) which blocks the BMP type I receptor site. As ERFE is trimeric and binds to dimeric BMP6 the complex retains a free LBD. Here, higher-order bundles could form using unpaired LBDs where a hexamer could bind to three BMP ligands in a terminal complex.

Comparing ERFE to other extracellular antagonists reveals interesting similarities and differences. BMP ligands bind type I receptors at the concave dimer interface and type II receptors on the convex surface.^42^ Figure 8 highlights how various inhibitors bind and occlude the type I binding site on the ligand. Noggin and Gremlin form extensive contacts on the knuckle region of the ligand and thread a peptide into the type I site (using a knob-in-hole).^24,26^ Both follistatin and ERFE bind using a more well-ordered helix to bind in the same pocket.^43^ Other non-inhibitory binding partners utilize helical motifs as well; ligand prodomains bind using an α1 helix and the repulsive guidance molecule (RGM) family, which act as critical BMP co-receptors, interact using a bundle of helices.^44,45^ Interestingly, the aforementioned helices and loops in BMP antagonists and binding partners thread through the type I receptor site in the opposite direction as the receptor helices. ERFE may separate itself from other characterized antagonistic interactions which occlude both type I and type II receptor sites, as our current data suggests ERFE only occludes type I receptor binding. Further studies may reveal additional points of contact between ligand and antagonist, or show that ERFE’s avidity allows it to circumvent the need for the extensive interactions seen in other inhibitors.

**Fig 8.**
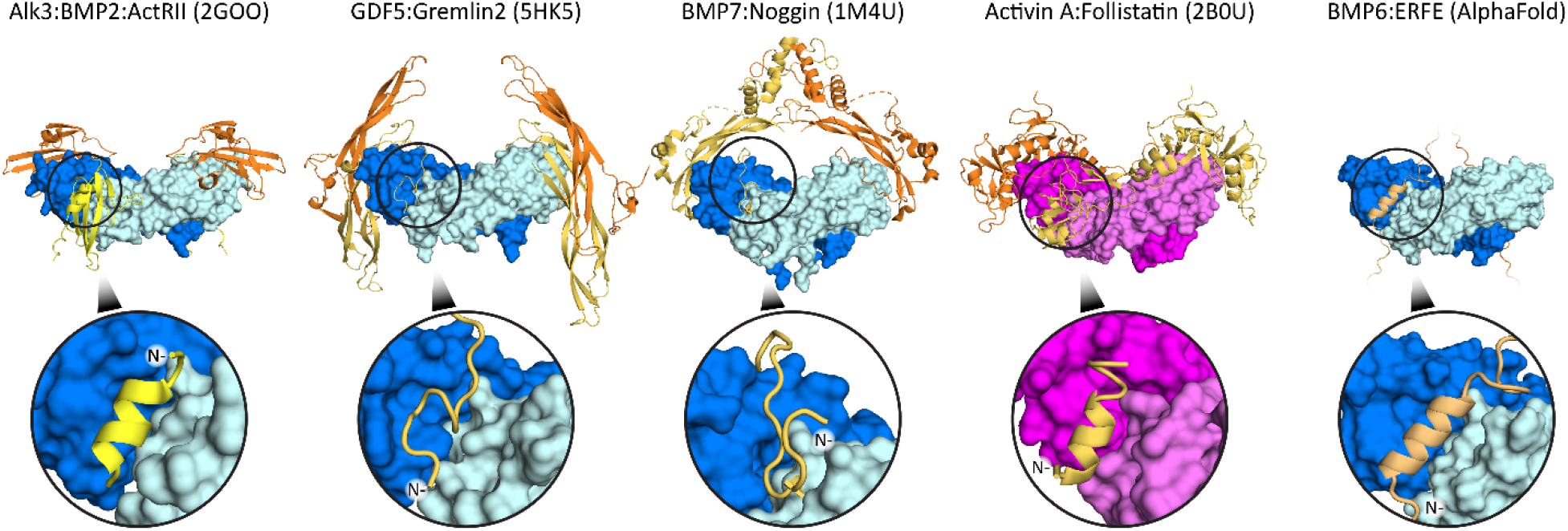
Structural comparison of TGF-β inhibitors. Extracellular antagonists utilize a similar strategy to block the type I binding site on the ligand. For reference, the ternary complex of BMP2 bound to the type I (Alk3) and type II (ActRII) receptor is shown on the left. Noggin and Gremlin2 both occlude BMP type I receptor binding with N-terminal unstructured loops that form ancillary interactions that complement the large buried surface area on the opposing side of the ligand. Follistatin interacts with the Activin A type I receptor pocket using an N-terminal helix. While ERFE is currently predicted to interact with the type I binding site in a manner similar to other inhibitors, our present model lacks additional interactions with other regions of the ligand. The N-terminus of each interacting segment is labeled to highlight that the inhibitors all run counter to the receptor.

In conclusion, we have demonstrated that the basic functional unit of ERFE is a trimer, as all of its oligomerizing features all stabilize trimer formation. Meanwhile, oligomerization above a trimer is dispensable for inhibition of BMP signaling. Finally, we identified a key conserved segment in ERFE’s unstructured domain that is critical for activity and occlusion of the type I receptor site. Whether this is the only molecular feature that interacts with BMP ligands is not known. Future structural studies are required to both validate the modeling efforts and determine the extent of the ERFE:BMP interaction. Furthermore, while our study also highlights the specificity of ERFE for certain BMP ligands, the molecular basis for this specificity remains undetermined.

## Materials and Methods

### Recombinant ligands and detection antibodies

Activin A, AMH, BMP2, and GDF5 were all produced as previously described.^46–48^ Mature GDF11 was a generous gift from Elevian. BMP6 was a generous gift from OSTEOGROW. ERFE was detected using DYKDDDDK Epitope Tag HRP-conjugated Antibody (R&D, Cat. # HAM85291) and FAM132B Polyclonal Antibody (Invitrogen, Cat. #PA5-67448). Recombinant ALK3-Fc fusion protein was purchased from R&D (Cat. No. 2406-BR-100).

### Protein production and purification

ERFE production was performed using a N-terminally flag tagged hERFE construct (residues 44-354) in the mammalian expression vector pcDNA3.1 generously given by Tomas Ganz (UCLA).^27^ Purified plasmid was transfected into Expi293 cells at density 2.0x10^6^ using polyethylenimine (PEI). Media was collected at day 3 applied directly to a HiPrep Heparin FF affinity column. Protein was eluted with a gradient from 500-650mM NaCl at pH7.5 with 20mM HEPES and 5mM CaCl2. A single peak containing ERFE was pooled and dialyzed into 20mM HEPES pH 7.5, 150mM NaCl, and 5mM CaCl_2_, concentrated to 1mg/mL, and either used immediately or flash frozen. WT yields ranged between 1-2 mg/L and varied mutant to mutant.

For ERFE mutants that were unable to be purified solely via heparin affinity, conditioned media was first applied to αFlag resin and eluted with 100mM Glycine pH 2.5. After neutralization with 500mM HEPES pH 7.5, it was applied to a heparin column and purified in the same manner described above. Yields for this preparatory method were slightly lower than exclusively using heparin affinity chromatography. No change in activity was seen when WT ERFE was purified using this method (Supporting figure **1.C**).

BMP7 was produced using a modified protocol from Cate **et al**.^**49**^ In summary, BMP7 cDNA in a pRK5 mammalian expression vector was transfected into Expi293 cells with PEI. Conditioned media was harvested 3 days post-transfection and applied to a HiTrap SP Sepharose cation exchange column at pH7.5 and eluted using a salt gradient (HEPES pH 7.5, 0-1M NaCl, 1mM EDTA). Fractions were pooled and dialyzed into 4mM HCl overnight before being loaded onto a reversed phase C18 column, eluted with an acetonitrile gradient and flash frozen after dialysis into 4mM HCl. Average yields were approximately 0.35mg mature ligand per liter conditioned media. Activity was confirmed via luciferase reporter assay in BRITER cells (Supporting figure **1.D**).^28^

### Luciferase reporter assay

A BMP-responsive luciferase reporter osteoblast cell line (BRITER, RRID:CVCL_0P40), provided by Amitabha Bandyopadhyay (Indian Institute of Technology), was used to measure BMP activity as previously reported.^28,50,51^ Briefly, cells were grown overnight in α-minimal essential medium with 10% (v/v) FBS and 100 µg/ml hygromycin B in a 96 well plate at 37°C in 5% CO2. The medium was replaced with DMEM and cells were starved for 5 h. Activity assays were performed by titrating ligands in DMEM, while inhibition assays were performed by titrating inhibitor and adding constant ligand. The media was replaced with ligand or ligand+inhibitor and incubated at the same conditions for 3 h before luminescence was read using a BioTek Synergy H1 plate reader. Inhibitory curves were normalized using ligand alone as 100% signal. Curve fits and both EC_50_ and IC_50_ values were generated using GraphPad Prism. All IC50 values were calculated using a variable hill slope. Each experiment consisted of 3 technical replicates and either 2 or 3 biological replicates were performed, as indicated.

### Analytical Ultracentrifugation

Protein samples were used post-dialysis and were never frozen. Sedimentation velocity experiments were run using Beckman Optima XL-I analytical ultracentrifuge (Beckman Coulter, Fullerton, CA), An60-Ti rotor, and absorbance optics. Samples at 0.5 mg/mL were loaded into Beckman AUC sample cells with 12-mm optical path two-channel centerpieces, with matched buffer in the reference sector. Full-length ERFE constructs were centrifugated at 20,000 rpm, while ΔC1Q mutants were centrifugated at 40,000 RPM, absorbance was measured at 230nm to maximize signal, and the experiment was performed at 20°C. SEDFIT was used to determine a preliminary frictional ratio using continuous c(s,ff0) distribution. The most commonly occurring frictional ratio was used as a starting value to solve for a precise frictional ratio using c(s) distributions, which yielded predicted molecular weights. In rare cases where solved c(s) and c(s,ff0) frictional ratios did not agree, the 3 most frequent c(s,ff0) frictional ratio were averaged, which ameliorated the issue. All data sets were normalized on a scale from 0 to 1, using the highest and lowest value.

### AlphaFold and Consurf

AlphaFold 2.3.0 was used to create models of the ERFE:BMP6 complex. Five AMBER relaxed models were generated. Analysis was focused on the top ranked model. Potential interactions were analyzed using PDBePISA^52^ and PyMOL.^53^ For analysis of evolutionary conservation, ERFE (AA 29-354) was analyzed using the ConSurf webserver with default settings.^30,31^ Results were used to examine conservation of predicted interacting residues before mutagenesis.

### Surface Plasmon Resonance (SPR)

Binding kinetics of BMP6 to Alk3 was determined by surface plasmon resonance using a BIAcore T-200 optical sensor system. 0.45ug/mL Alk3-Fc was first captured on a protein a chip (Cat. # 29127555), establishing a baseline of 85 response units (RU). 10nM constant BMP6 with ERFE concentrations serially diluted in SPR running buffer (20 mM HEPES pH 7.4, 350 mM NaCl, 0.005 % P-20, 0.5 mg/mL BSA, 5 mM EDTA) to 80–2.5 nM, were flowed over the chip at 50 μL/min for 300 s to observe association, then washed off for 900 s to observe dissociation. Additionally, ERFE alone at 80nM was flowed over the chip as a control. Regeneration was performed using 10mM glycine pH 1.8.

## Acknowledgements and author contributions

JM and TBT planned the experiments. JM performed the research and analyzed data with TBT. EL assisted with luciferase assay experiments. JM and TBT wrote the manuscript. Special thanks to the TT lab for assisting with data interpretation and experimental design

## References

1. Kautz, L. et al. Identification of erythroferrone as an erythroid regulator of iron metabolism. Nat. Genet. 46, 678–684 (2014).

2. Pigeon, C. et al. A New Mouse Liver-specific Gene, Encoding a Protein Homologous to Human Antimicrobial Peptide Hepcidin, Is Overexpressed during Iron Overload*. J. Biol. Chem. 276, 7811–7819 (2001).

3. Andriopoulos, B. et al. BMP-6 is a key endogenous regulator of hepcidin expression and iron metabolism. Nat. Genet. 41, 482–487 (2009).

4. Truksa, J., Peng, H., Lee, P. & Beutler, E. Bone morphogenetic proteins 2, 4, and 9 stimulate murine hepcidin 1 expression independently of Hfe, transferrin receptor 2 (Tfr2), and IL-6. Proc. Natl. Acad. Sci. 103, 10289–10293 (2006).

5. Andolfo, I. et al. The BMP-SMAD pathway mediates the impaired hepatic iron metabolism associated with the ERFE-A260S variant. Am. J. Hematol. 94, 1227–1235 (2019).

6. Wang, C.-Y. et al. Erythroferrone lowers hepcidin by sequestering BMP2/6 heterodimer from binding to the BMP type I receptor ALK3. Blood 135, 453–456 (2020).

7. Kautz, L. et al. Erythroferrone contributes to hepcidin suppression and iron overload in a mouse model of β-thalassemia. Blood 126, 2031–2037 (2015).

8. Arezes, J. et al. Antibodies against the erythroferrone N-terminal domain prevent hepcidin suppression and ameliorate murine thalassemia. Blood 135, 547–557 (2020).

9. Gardenghi, S. et al. Ineffective erythropoiesis in β-thalassemia is characterized by increased iron absorption mediated by down-regulation of hepcidin and up-regulation of ferroportin. Blood 109, 5027– 5035 (2007).

10. Schäffler, A. & Buechler, C. CTRP family: linking immunity to metabolism. Trends Endocrinol. Metab. TEM 23, 194–204 (2012).

11. Ressl, S. et al. Structures of C1q-like Proteins Reveal Unique Features among the C1q/TNF Superfamily. Structure 23, 688–699 (2015).

12. Païdassi, H. et al. C1q Binds Phosphatidylserine and Likely Acts as a Multiligand-Bridging Molecule in Apoptotic Cell Recognition1. J. Immunol. 180, 2329–2338 (2008).

13. Pascolutti, R., Erlandson, S. C., Burri, D. J., Zheng, S. & Kruse, A. C. Mapping and engineering the interaction between adiponectin and T-cadherin. J. Biol. Chem. 295, 2749–2759 (2020).

14. Berisio, R., Vitagliano, L., Mazzarella, L. & Zagari, A. Crystal structure of the collagen triple helix model [(Pro-Pro-Gly)10]3. Protein Sci. 11, 262–270 (2002).

15. Erlich, P. et al. Complement Protein C1q Forms a Complex with Cytotoxic Prion Protein Oligomers. J. Biol. Chem. 285, 19267–19276 (2010).

16. Wong, G. W. et al. Identification and characterization of CTRP9, a novel secreted glycoprotein, from adipose tissue that reduces serum glucose in mice and forms heterotrimers with adiponectin. FASEB J. 23, 241–258 (2009).

17. Wong, G. W. et al. Molecular, biochemical and functional characterizations of C1q/TNF family members: adipose-tissue-selective expression patterns, regulation by PPAR-γ agonist, cysteine-mediated oligomerizations, combinatorial associations and metabolic functions. Biochem. J. 416, 161–177 (2008).

18. Briggs, D. B. et al. Disulfide-Dependent Self-Assembly of Adiponectin Octadecamers from Trimers and Presence of Stable Octadecameric Adiponectin Lacking Disulfide Bonds in Vitro. Biochemistry 48, 12345–12357 (2009).

19. Seldin, M. M., Peterson, J. M., Byerly, M. S., Wei, Z. & Wong, G. W. Myonectin (CTRP15), a Novel Myokine That Links Skeletal Muscle to Systemic Lipid Homeostasis. J. Biol. Chem. 287, 11968–11980 (2012).

20. Stewart, A. N., Little, H. C., Clark, D. J., Zhang, H. & Wong, G. W. Protein Modifications Critical for Myonectin/Erythroferrone Secretion and Oligomer Assembly. Biochemistry 59, 2684–2697 (2020).

21. Gipson, G. R. et al. Structural perspective of BMP ligands and signaling. Bone 140, 115549 (2020).

22. Hinck, A. P., Mueller, T. D. & Springer, T. A. Structural Biology and Evolution of the TGF-β Family. Cold Spring Harb. Perspect. Biol. 8, a022103 (2016).

23. Kattamuri, C. et al. Members of the DAN Family Are BMP Antagonists That Form Highly Stable Noncovalent Dimers. J. Mol. Biol. 424, 313–327 (2012).

24. Nolan, K. et al. Structure of Gremlin-2 in Complex with GDF5 Gives Insight into DAN-Family-Mediated BMP Antagonism. Cell Rep. 16, 2077–2086 (2016).

25. Troilo, H. et al. Nanoscale structure of the BMP antagonist chordin supports cooperative BMP binding. Proc. Natl. Acad. Sci. 111, 13063–13068 (2014).

26. Groppe, J. et al. Structural basis of BMP signalling inhibition by the cystine knot protein Noggin. Nature 420, 636–642 (2002).

27. Ganz, T. et al. Immunoassay for human serum erythroferrone. Blood 130, 1243–1246 (2017). 28.

28. Yadav, P. S., Prashar, P. & Bandyopadhyay, A. BRITER: A BMP Responsive Osteoblast Reporter Cell Line. PLOS ONE 7, e37134 (2012).

29. Melchert, J. et al. The secreted BMP antagonist ERFE is required for the development of a functional circulatory system in Xenopus. Dev. Biol. 459, 138–148 (2020).

30. Ashkenazy, H. et al. ConSurf 2016: an improved methodology to estimate and visualize evolutionary conservation in macromolecules. Nucleic Acids Res. 44, W344–W350 (2016).

31. Yariv, B. et al. Using evolutionary data to make sense of macromolecules with a”face-lifted” ConSurf. Protein Sci. 32, e4582 (2023).

32. Jumper, J. et al. Highly accurate protein structure prediction with AlphaFold. Nature 596, 583– 589 (2021).

33. [Preprint] Evans, R. et al. Protein complex prediction with AlphaFold-Multimer. 2021.10.04.463034 Preprint at 10.1101/2021.10.04.463034 (2021).

34. Keller, S., Nickel, J., Zhang, J.-L., Sebald, W. & Mueller, T. D. Molecular recognition of BMP-2 and BMP receptor IA. Nat. Struct. Mol. Biol. 11, 481–488 (2004).

35. Kirsch, T., Sebald, W. & Dreyer, M. K. Crystal structure of the BMP-2–BRIA ectodomain complex. Nat. Struct. Biol. 7, 492–496 (2000).

36. Saremba, S. et al. Type I receptor binding of bone morphogenetic protein 6 is dependent on Nglycosylation of the ligand. FEBS J. 275, 172–183 (2008).

37. Isaacs, M. J. et al. Bone Morphogenetic Protein-2 and -6 Heterodimer Illustrates the Nature of Ligand-Receptor Assembly. Mol. Endocrinol. 24, 1469–1477 (2010).

38. Castro-Mollo, M. et al. The hepcidin regulator erythroferrone is a new member of the erythropoiesis-iron-bone circuitry. eLife 10, e68217 (2021).

39. Li, K. et al. Myonectin Predicts the Development of Type 2 Diabetes. J. Clin. Endocrinol. Metab. 103, 139–147 (2018).

40. Little, H. C. et al. Myonectin deletion promotes adipose fat storage and reduces liver steatosis. FASEB J. 33, 8666–8687 (2019).

41. Persikov, A. V., Ramshaw, J. A. M. & Brodsky, B. Prediction of Collagen Stability from Amino Acid Sequence*. J. Biol. Chem. 280, 19343–19349 (2005).

42. Allendorph, G. P., Vale, W. W. & Choe, S. Structure of the ternary signaling complex of a TGF-β superfamily member. Proc. Natl. Acad. Sci. 103, 7643–7648 (2006).

43. Thompson, T. B., Lerch, T. F., Cook, R. W., Woodruff, T. K. & Jardetzky, T. S. The Structure of the Follistatin:Activin Complex Reveals Antagonism of Both Type I and Type II Receptor Binding. Dev. Cell 9, 535–543 (2005).

44. Cotton, T. R. et al. Structure of the human myostatin precursor and determinants of growth factor latency. EMBO J. 37, 367–383 (2018).

45. Malinauskas, T., Peer, T. V., Bishop, B., Mueller, T. D. & Siebold, C. Repulsive guidance molecules lock growth differentiation factor 5 in an inhibitory complex. Proc. Natl. Acad. Sci. 117, 15620–15631 (2020).

46. Hart, K. N. et al. Structure of AMH bound to AMHR2 provides insight into a unique signaling pair in the TGF-β family. Proc. Natl. Acad. Sci. 118, e2104809118 (2021).

47. Gipson, G. R. et al. Formation and characterization of BMP2/GDF5 and BMP4/GDF5 heterodimers. BMC Biol. 21, 16 (2023).

48. Goebel, E. J. et al. Structural characterization of an activin class ternary receptor complex reveals a third paradigm for receptor specificity. Proc. Natl. Acad. Sci. 116, 15505–15513 (2019).

49. Cate, R. L. et al. The anti-Müllerian hormone prodomain is displaced from the hormone/prodomain complex upon bivalent binding to the hormone receptor. J. Biol. Chem. 298, 101429 (2021).

50. Gipson, G. R., Kattamuri, C., Czepnik, M. & Thompson, T. B. Characterization of the different oligomeric states of the DAN family antagonists SOSTDC1 and SOST. Biochem. J. 477, 3167–3182 (2020).

51. Nolan, K. et al. Structure of Neuroblastoma Suppressor of Tumorigenicity 1 (NBL1): INSIGHTS FOR THE FUNCTIONAL VARIABILITY ACROSS BONE MORPHOGENETIC PROTEIN (BMP) ANTAGONISTS*. J. Biol. Chem. 290, 4759–4771 (2015).

52. Krissinel, E. & Henrick, K. Inference of macromolecular assemblies from crystalline state. J. Mol. Biol. 372, 774–797 (2007).

53. Schrödinger, LLC. The PyMOL Molecular Graphics System, Version 1.8. (2015).

54. Varadi, M. et al. AlphaFold Protein Structure Database: massively expanding the structural coverage of protein-sequence space with high-accuracy models. Nucleic Acids Res. 50, D439–D444 (2022).

