## Supplemental Figures 1-3 for "Characterization of erythroferrone oligomerization and its impact on BMP antagonism"

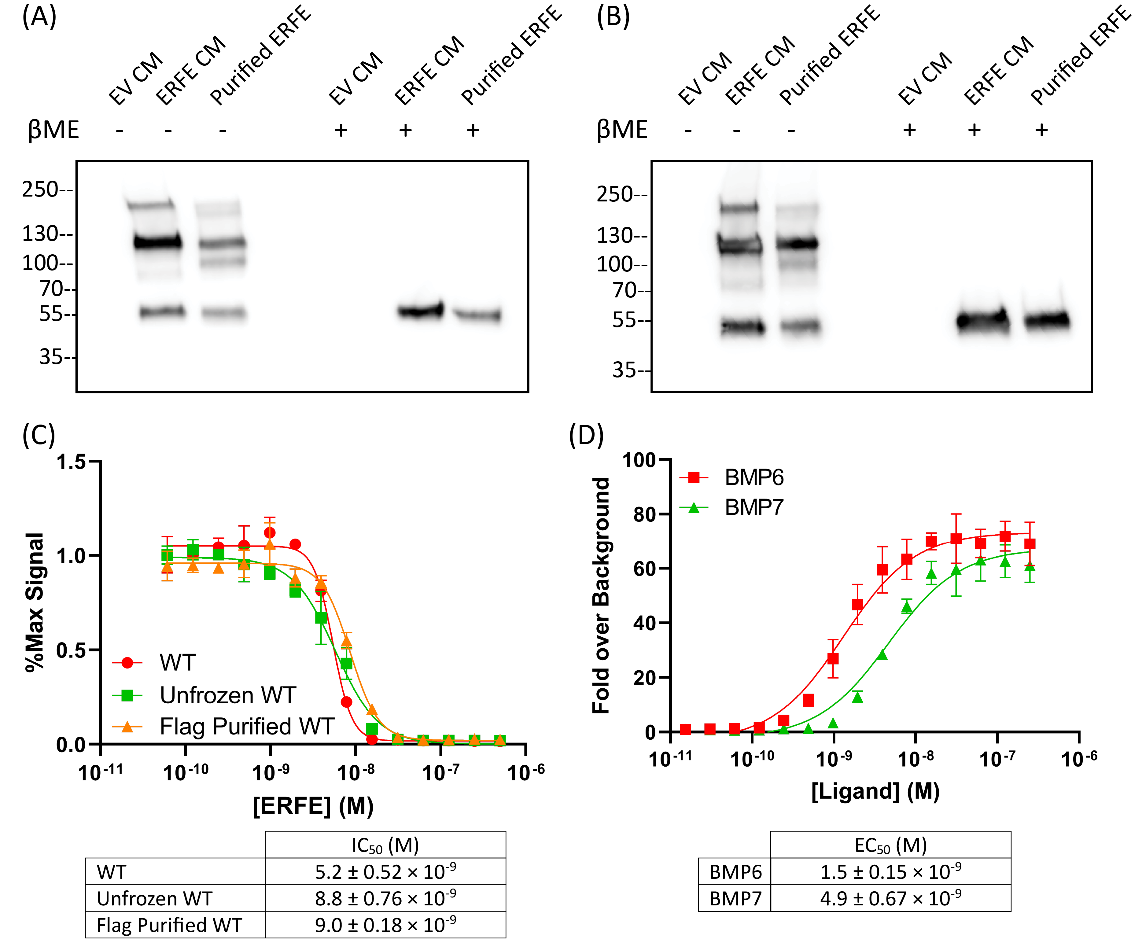

Supporting Figure 1: **Verification of Reagents and Purification Methods**. Purified ERFE was analyzed via western blot probing with (**A**) αERFE or (**B**) αFLAG-tag antibodies. (**C**) WT ERFE did not change its potency regardless of freezing or purifying via flag affinity followed by heparin affinity chromatography. (**D**) BMP6 and BMP7 used in this paper are potent activators of BMP signaling in BRITER cells. Statistical analysis is detailed in materials and methods, mean ± of 3 biological replicates shown. SEM SD of 3 points within a single technical replicate shown with error bars. (n=3, representative curve shown).

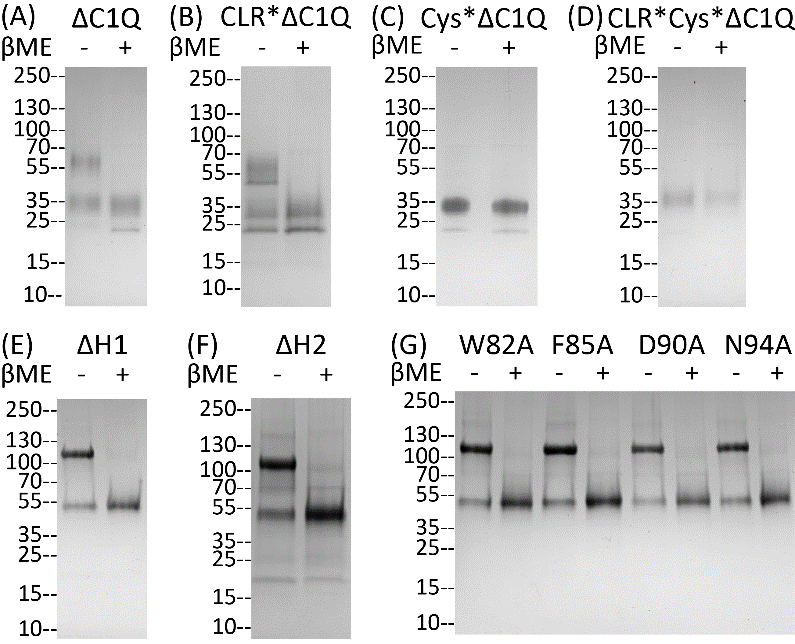

Supporting Figure 2: **Verification of ERFE Mutant Purity**. (**A-D**) To verify protein quality and purity, 5ug of the specified ΔC1Q mutant was analyzed by SDS-PAGE and stained with coomassie blue. (**E-G**) The same was done for ΔH1, ΔH2, and single point mutations in the LBD of ERFE.

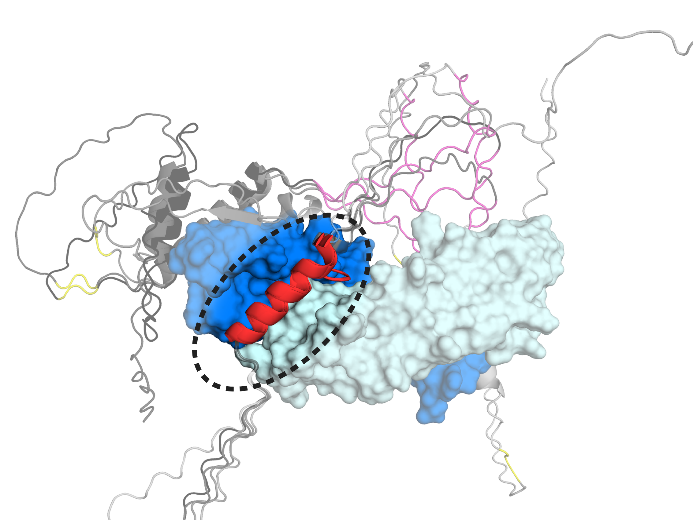

Supporting Figure 3: **Analysis of ERFE Helix Predicted to Interact with BMP6.** (**A**) AlphaFold was used to generate 5 AMBER relaxed models, all of which were overlayed. H1 (red) was the only feature that was consistently placed in the same position on the BMP ligand. The cysteines (yellow), CLR (pink), and H2 (black/grey) were never consistently placed.
